# Long- and short-term history effects in a spiking network model of statistical learning

**DOI:** 10.1101/2021.09.22.461372

**Authors:** Amadeus Maes, Mauricio Barahona, Claudia Clopath

## Abstract

The statistical structure of the environment is often important when making decisions. There are multiple theories of how the brain represents statistical structure. One such theory states that neural activity spontaneously samples from probability distributions. In other words, the network spends more time in states which encode high-probability stimuli. Existing spiking network models implementing sampling lack the ability to learn the statistical structure from observed stimuli and instead often hard-code a dynamics. Here, we focus on how arbitrary prior knowledge about the external world can both be learned and spontaneously recollected. We present a model based upon learning the inverse of the cumulative distribution function. Learning is entirely unsupervised using biophysical neurons and biologically plausible learning rules. We show how this prior knowledge can then be accessed to compute expectations and signal surprise in downstream networks. Sensory history effects emerge from the model as a consequence of ongoing learning.

## INTRODUCTION

There is an ever-increasing body of evidence indicating that the brain takes the statistical regularities in the environment into account (Barlow 1961; Ruderman and Bialek 1994; Maye et al. 2002). It might do so in a number of ways. Firstly, *a priori* knowledge of sensory stimuli influences perception. For example, an expected stimulus might be encoded faster than an unexpected stimulus (Kok et al. 2017; Mazzucato et al. 2019), conversely, low-probability stimuli might trigger a stronger response to signal novelty or surprise (Ulanovsky et al. 2003; Khouri and Nelken 2015; Hamm et al. 2021). Secondly, knowledge of the statistics of a relevant variable influences decisions, potentially leading to biases (Akrami et al. 2018; Zylberberg et al. 2018; Lieder et al. 2019; Hachen et al. 2020; Meirhaeghe et al. 2021). For the brain to take prior statistical structure into account, it needs a way to learn and recollect such structure.

One line of investigation studies the neural ensemble as a building block for processing and computations in the brain, in line with Hebb’s postulate (Hebb 1949). Stimuli that are repeatedly presented, could be encoded in groups of neurons and be spontaneously replayed in the absence of the stimuli. Recently, experimental studies have started to explore these ideas in detail. It was found that neural ensembles are coactive transiently both during evoked activity and spontaneous activity (Luczak et al. 2009) and can be developed by repeated stimulation (Miller et al. 2014; Carrillo-Reid et al. 2016). Additionally, such neural ensembles can affect behaviour, elucidating their functional relevance (Carrillo-Reid et al. 2019; Carrillo-Reid 2021). In parallel, computational work has studied the plasticity mechanisms needed to develop neural ensembles in networks by stimulating the network repeatedly with a set of stimuli (Clopath et al. 2010; Litwin-Kumar and Doiron 2014; Zenke et al. 2015; Maes et al. 2020). The connectivity pattern which emerges from repeated stimulations is clustered, i.e. the excitatory neurons are strongly recurrently connected within the same cluster, and weakly connected between clusters (Peron et al. 2020). Such connectivity reverberates the activity within the same cluster and leads to random switching dynamics, i.e. the clusters switch between high and low activity states at random (Litwin-Kumar and Doiron 2012; Schaub et al. 2015). While spontaneous reactivations can be interpreted as a recollection of the previously applied stimuli, they do not depend on the probability by which the stimuli were applied. Hence, current models fail to incorporate the statistical structure of the stimuli.

Here, we propose a way in which the spontaneous activity of the network depends on the probabilities of the stimuli exposure, i.e. the network activity samples from the prior stimulus distribution. Specifically, we implement inverse transform sampling in the model and learn by repeatedly applying stimuli to the model, using biophysically realistic neurons and plasticity mechanisms. We then explore how this representation can be useful for computations. Firstly, sampling lends itself to performing Monte Carlo-type calculations. By reading out and integrating samples we show it is easy to compute expectations over functions. A specific example where the brain might compute expectations is when doing perceptual decision-making. In this context, the model exhibits long- and short-term history effects. These history effects originate from the plasticity in the model, slowly forgetting old stimuli and biasing decisions on a short time scale. Finally, we show how we can transform the representation into a more instantaneous code, potentially relevant for fast sensory processing and predictive coding.

## RESULTS

### A model designed to learn statistical structure

We design spiking networks to perform inverse transform sampling. Consider the random variable *X*, the cumulative distribution function *F*(*x*) and probability density function 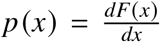. A sample *x* can be drawn from *p*(*x*), by the following textbook procedure: 1) take a sample from the uniform distribution 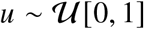; 2) transform the sample using the inverse of the cumulative distribution function, i.e. *x* = *F*^−1^(*u*). We first show that this procedure can be implemented using two spiking networks (Fig 1A). One network samples from the uniform distribution, using random dynamics. We call this first network the uniform sampler network. The second network represents the random variable and is driven by the first network. The weights from the first to the second network corresponds to the inverse of the cumulative distribution function and change using simple biologically plausible learning rules (Fig 1B). We call the second network the sensory network.

**Fig. 1.**
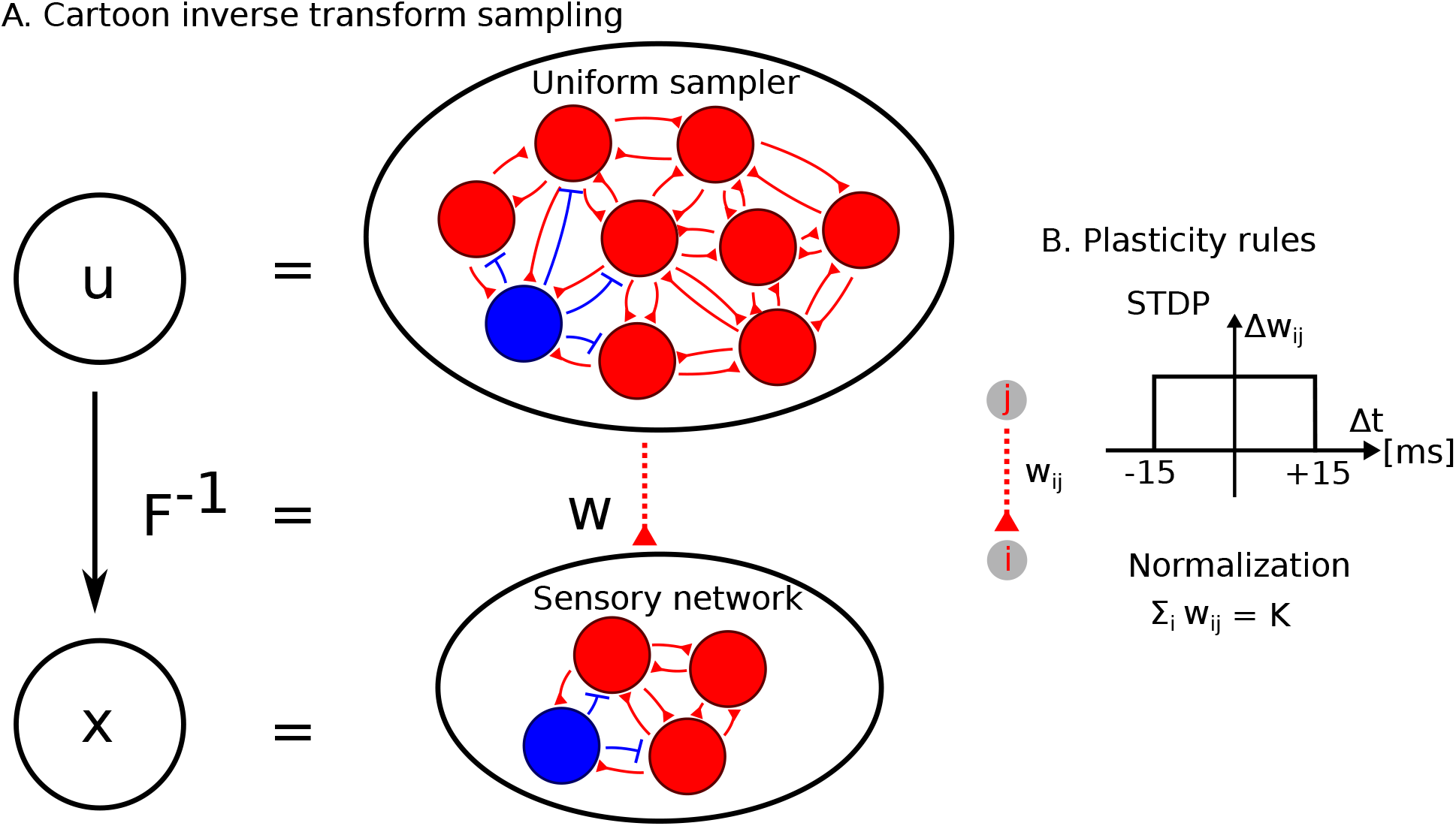
A model designed to learn statistical structure. (A) Cartoon of the model. Inverse transform sampling is mapped to two spiking networks. Uniform samples (top) are transformed through weights (middle) to facilitate sampling from the variable x (bottom). (B) There are two plasticity mechanisms, STDP provides potentiation and normalization provides depression (see Methods for details).

The uniform sampler network consists of excitatory and inhibitory neurons. We group the excitatory neurons in *C* disjoint clusters. Excitatory neurons are strongly connected to other excitatory neurons in the same cluster and weakly connected to all other excitatory neurons. The inhibitory neurons act as a single stabilizing pool. This connectivity structure leads to transiently active clusters, where each cluster activated at random silences the other clusters through the lateral inhibition (Schaub et al. 2015). Here, we fix this connectivity structure, but previous work has shown that such a structure can be learned using biologically plausible rules (Clopath et al. 2010; Ko et al. 2013; Litwin-Kumar and Doiron 2014; Maes et al. 2020). The random switching dynamics in this network can be interpreted as sampling from the uniform distribution, where the amount of probability in a single cluster amounts to 1/*C*. The sensory network encodes the external variable. The network is organized in the same way as the uniform sampler network. The number of clusters in the sensory network is 8 throughout the paper. This means that the sensory network discretizes the external variable of interest in 8 intervals. While the activity in the sensory network reverberates due to the recurrent clustered connectivity, the input from the uniform sampler controls the switches between sensory clusters. To summarize, the architecture leads to two approximations: 1) discretization of the uniform space in parts of 1|*C*; 2) discretization of the encoded external variable.

To train the model, we present samples of the external variable *X* sequentially. At each observation, external current activates the cluster in the sensory network encoding the observed sample. The first plasticity mechanism is a potentiation through spike-time-dependent plasticity (STDP). The cluster in the uniform sampler that happens to be active at the moment of the observation will potentiate its connections to the cluster in the sensory network. In this way, the model attributes an amount of 1/*C* probability to the observation. The second plasticity mechanism is a normalization, leading to the depression of the connections from the active cluster in the uniform sampler to non-active clusters in the sensor network. This mechanism ensures that each cluster in the uniform sampler projects only to a single cluster in the sensory network. In summary, the potentiation attributes an amount of probability to the new observation and the normalization removes the same amount of probability from an older observation. Repeated observations will shape the weights from the uniform sampler network to the sensory network, approximating the inverse of the cumulative distribution function. Different versions of both the STDP rule and normalization are commonly used to model synaptic plasticity (Zenke et al. 2013; Toyoizumi et al. 2014; Litwin-Kumar and Doiron 2014; Zenke et al. 2015).

### The model learns through repeated observations

We first show how the inverse transform *F*^−1^(*u*) can be learned and analyze the accuracy. Learning is unsupervised: samples from the target distribution are observed by interacting with the external world (Fig 2A). We assume that the network has already learned a previous distribution *p*(*x*). However, to emphasize the learning abilities of the model, we now show a new target distribution (Fig 2B). Samples from the target distribution are observed at a rate of 5 Hz so that every 200 ms we apply an external input to the sensory network (Fig S1A). The plastic weights projecting from the uniform sampler network to the sensory network will change to reflect the inverse of the cumulative distribution function of the target distribution (Fig S1B). We obtain learning curves by taking the L1 error between the normalized weight matrix and the target inverse transform (see Methods) (Fig 2C). Learning is faster when there are fewer clusters in the uniform sampler network, however, there are larger fluctuations in the error (Fig 2D). This trade-off is a result of the discretization of the uniform distribution scaling as 1/*C*. We conclude that the model is able to learn the inverse transform by repeated observations of the variable.

**Fig. 2.**
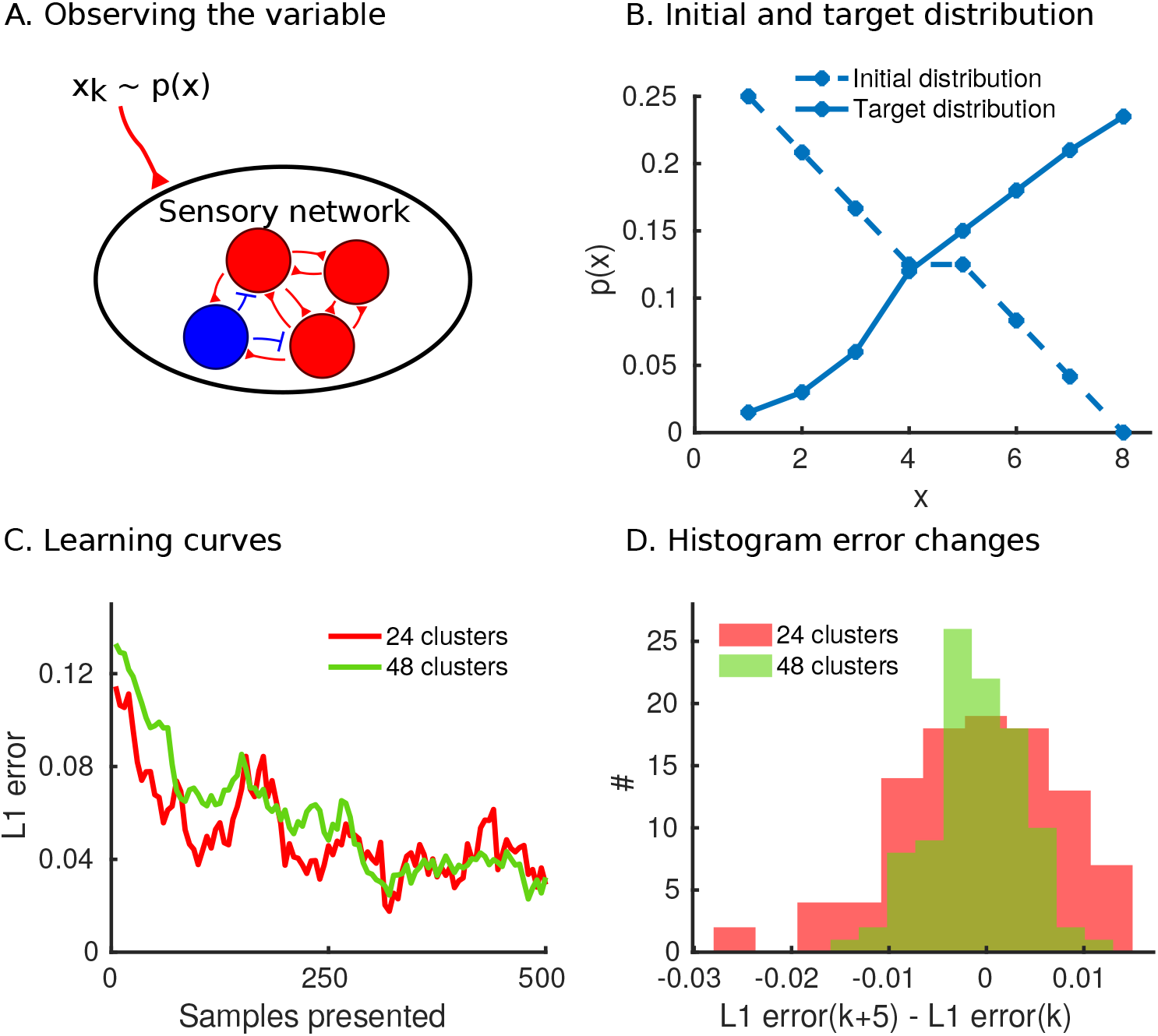
The model learns through repeated observations. (A) The sensory network receives external input from observing samples *x_k_* of the variable *X*. (B) An initial distribution is encoded in the weights, and samples from the target distribution are presented. (C) Learning curves for a uniform sampler with 24 (red) and 48 (green) clusters. The L1 error is computed between the normalized weight matrix and the target distribution. (D) The change in error is measured after every fifth sample presentation and the resulting values plotted in a histogram. The error fluctuations are higher for a lower number of clusters in the uniform sampler network (Mann-Whitney U-test on the absolute values, *p* < 10^−4^).

### The model performs sampling during spontaneous dynamics

We then verified the sampling behaviour of the model. From the theory, we expect the sensory network to sample from the target distribution. Simulations of spontaneous dynamics of the model show that, over a sufficiently long time, the clusters in the uniform sampler network are activated uniformly. Additionally, activations of clusters in the sensory network with a higher target probability are more likely to occur (Fig 3A). Quantitatively, we can measure the KL-divergence between the neural activity and the target distribution as a function of time (Fig 3B). We construct an empirical distribution from the neural activity by counting the fraction of time that each cluster is active (see Methods for details). The KL-divergence decreases with increasing sampling time, indicating a time frame of seconds to obtain an empirical distribution close to the target distribution. The sampling behaviour of the model is close to the behaviour of a random number generator. Samples drawn from the target distribution, using a random number generator at 8 Hz, i.e. at about the switching rate in the uniform sampler, yield very similar KL-divergence curves. To conclude, we show that in practice the neural activity of the sensory network approximately samples from the target distribution.

**Fig. 3.**
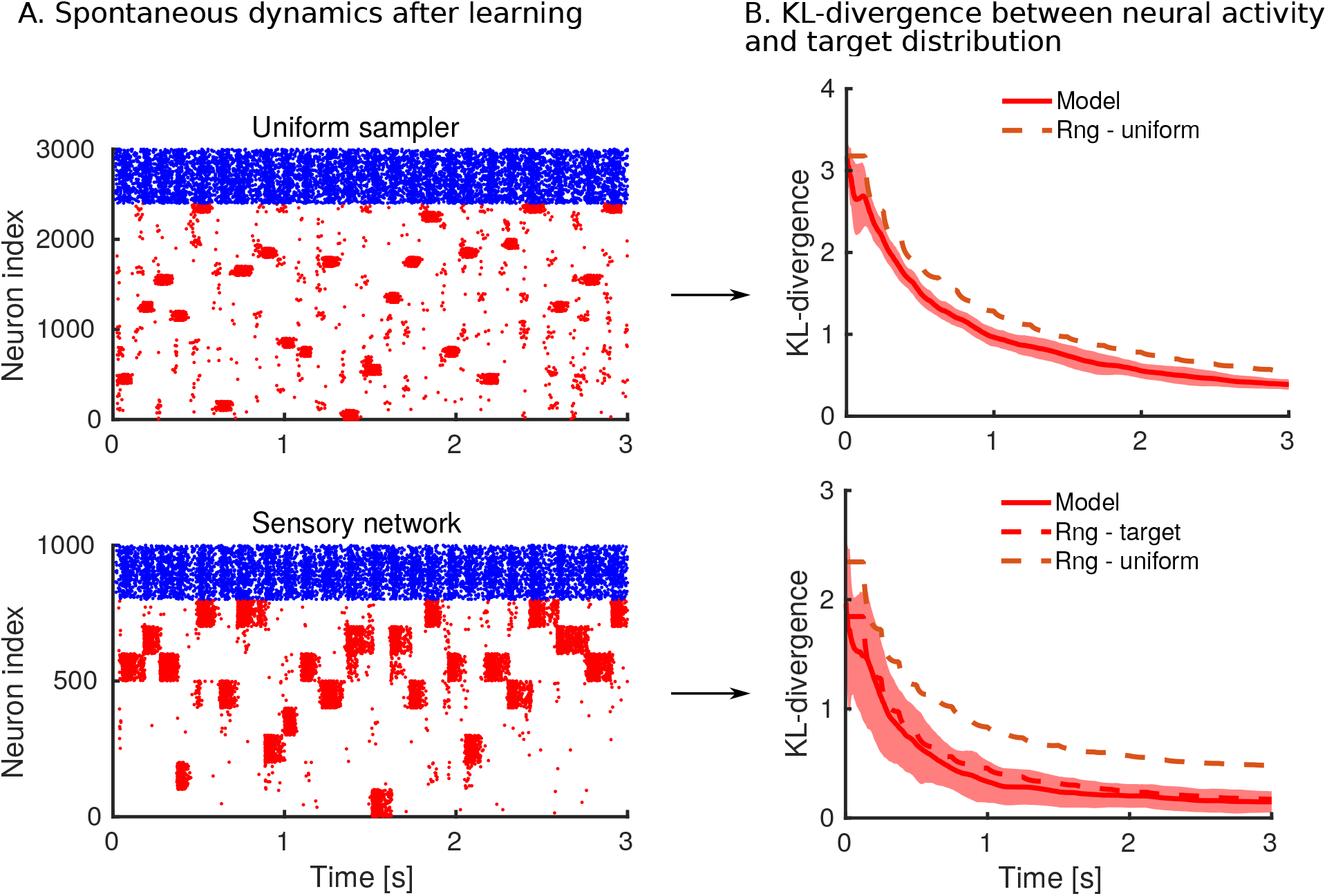
The model performs sampling during spontaneous dynamics. (A) Spike raster of the uniform sampler network (top) and sensory network (bottom). Red dots are spikes of excitatory neurons and blue dots are spikes of inhibitory neurons. The spike raster shows that the clusters in the uniform sampler switch randomly (at about ~ 8 Hz), and the clusters in the sensory network are active according to the stored distribution. (B) We measure the KL-divergence. The KL-divergence at time *t* takes into account all the neural activity from zero seconds to *t* seconds (see Methods for details). The red full lines are the KL-divergence between neural activity in the uniform sampler and the uniform distribution (top) and the KL-divergence between neural activity and target distribution of Fig 2B (bottom). The shaded area indicates one standard deviation from the mean (25 simulations). The red and brown dashed lines show the mean of the KL-divergence over 100 simulations when using a random number generator (rng) to draw samples from the target distribution and uniform distribution respectively, rather than using the model to generate samples.

### The model can provide samples for the computation of expectations

We just showed the ability of the model to learn the inverse transform and sample from the target distribution. We next wondered how this representation can be useful for downstream computations. A natural first idea is to use these samples to compute expectations of functions. Specifically, we can implement simple Monte Carlo approximations to compute integrals of the following type:

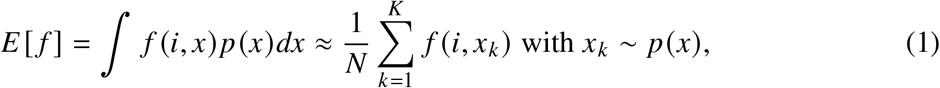

where *f*(*i, x*) is a function of the variable *X* and other inputs *i*. Because the samples *x_k_* become available over time, there has to be an integration mechanism updating the expectation over time. Defining *r_t_* to be the integration variable at time *t* which approximates the expectation *E*[*f*], we implement the integration as follows: 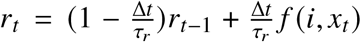 (forward Euler). *x_t_* is the sample produced by the model at time *t*, Δ*t* is the simulation time step and *τ_r_* is the time constant of integration. This way of computing expectations is modular and flexible compared to a system that integrates the sampling and function in one network. Expectations using arbitrary distributions may be computed in this way, where the distributions can be relearned while the function remains unchanged (Fig 4A).

**Fig. 4.**
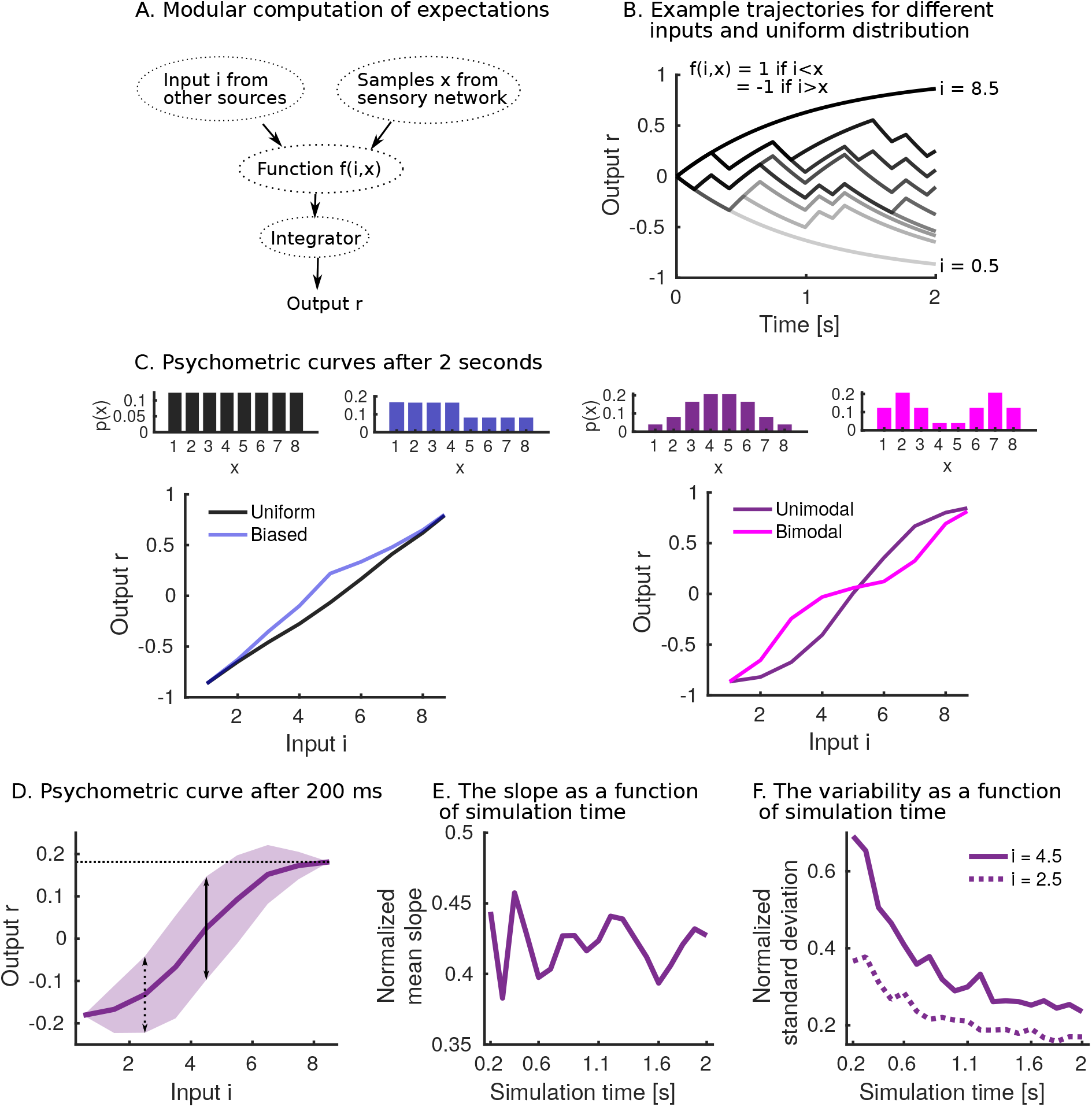
The model can provide samples for the computation of expectations. (A) Cartoon of computation, samples are internally generated by the inverse transform model and read-out by a downstream network which computes a function. This function is then integrated and generates an approximation of the expectation. (B) An input *i* is given, *i* = 0.5 to *i* = 8.5 in steps of one, and for each input the output is computed. (C) Psychometric curves after 2 seconds of simulating the system for varying distributions (averaged over 20 such simulations). (D) Psychometric curve after 200 ms of simulating the system, for the unimodal distribution. The shaded area is one standard deviation from the mean on each side (100 simulations). The horizontal dotted line indicates the maximum output at *i* = 8.5 (used for normalization). The dotted and full vertical lines indicate the standard deviations at *i* = 2.5 and *i* = 4.5 respectively. (E) The normalized mean of the slope as a function of simulation time remains constant (see Methods). (F) The normalized standard deviation of the output reduces with simulation time.

As an example, we consider the following indicator function: *f*(*i, x*) = 1 if *i* > *x* and *f*(*i, x*) = −1 if *i* < *x*. This function may be relevant to perceptual decision-making involving a binary choice.

There are two categories (+1/−1), and a choice between the categories is made based on a stimulus *i*. It is as of yet unclear how such decisions are made. One way in which a decision could be made is by taking the statistics of the stimulus *p*(*x*) into account rather than a decision boundary (Hachen et al. 2020). We simulate a decision by drawing an input *i* (*i* = [0.5, 8.5]) and computing and integrating *f* (*i, x_t_*). First, we simulate the system for 2 seconds for varying inputs and distributions (Fig 4B) and obtain psychometric curves by plotting the output *r* after 2 seconds as a function of the input *i* (Fig 4C). We observe clear differences in the psychometric curves, as a consequence of the different distributions *p*(*x*). Indeed, the output *r* is proportional to how likely the input *i* is larger than the random variable *X*: 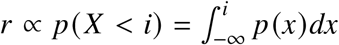. We first compare the uniform distribution with a biased distribution. The biased distribution has more probability mass in the interval *x* = [1,4] than in the interval *x* = [5,8], leading to an upwards biased psychometric curve compared to the uniform distribution. Then, we compare a unimodal and bimodal distribution. Here, the different distributions lead to a notable difference in the slopes of the psychometric curve. These theoretical psychometric curves can act as a prediction for future perceptual decision-making studies where the stimulus distribution is varied. Note, however, that we remain agnostic to the mechanism of the actual decision for one of the two categories. If the probability to choose category +1 is a monotonously increasing function of the output *r*, then the experimentally observed psychometric curves would be the result of transforming our theoretical psychometric curves by that function.

We wondered next to what extent the decision time affects the psychometric curve. The shape of the psychometric curve is not dependent on the amount of decision time available, as long as the curve is averaged over many individual decisions (Fig 4D-E). The sampling mechanism explains the independence of the shape on decision time. The samples are drawn independently, and an output *r* generated after a long decision time is equal to an average over multiple outputs *r* generated by short decision times. Unlike the average shape, the variability around the psychometric curve is affected by the decision time (Fig S2). When normalized, the variability reduces strongly with decision time (Fig 4F). Moreover, we also show that a “simpler” input has a lower variability than a “more difficult” input. An input is “simpler” when it is further away from the mean of the distribution, in which case it is easier to classify the input into one of the two categories. The inverse relationship between decision time and variability means we need more data to make a good estimation of the choice behaviour for short decision times. We conclude that the sampling mechanism can be useful for downstream networks in the context of computing expectations. Specifically, we predict differences between psychometric curves which are dependent on the stimulus distributions. The psychometric curves do not depend on the decision time but require more data to accurately estimate when decision times are shorter.

### The model exhibits long- and short-term history effects

We next wondered whether the model exhibits history effects. Recent work has shown sensory history-dependent biases in decision-making on both short (a few trials) and long time scales (~100 trials) (Mochol et al. 2021; Tervo et al. 2021). We expect history effects in the model, because the plastic weights adapt to newly observed samples, redistributing the probability mass continually (Fig S3). When we switch between target distributions, it takes about 100 samples to forget the old target distribution entirely (Fig 5A). The psychometric curve, measured shortly after the switch takes place, is an interpolation of the psychometric curves of the old and new target distributions (Fig 5B). On a shorter time scale, we look at the effect of the last five observed stimuli on the output, given input *i* = 4.5 (Fig 5C), when the same target distribution is presented (steady-state). We use here the bimodal target distribution to test for a short-term history effect. When regressing the mean of the last five samples on the normalized output, we observe a significant effect (Fig 5D). The short-term history can bias the output up to around 5%. This is an attractive bias, in the sense that the short-term mean of stimuli pulls the mean of the stored distribution in the model towards it. The bias arises in the model because the short-term statistics of stimuli can substantially differ from the target distribution. Such attractive biases are observed empirically at different strengths and in a wide variety of tasks, from delayed comparison tasks (Akrami et al. 2018), to the categorization of sounds (Chambers et al. 2017) and rating the attractiveness of faces (Xia et al. 2016). When the plasticity in the model is frozen, the short-term history effect becomes insignificant, further confirming the bias emerges due to learning of the stimulus distribution (Fig 5E). Other contributions to choice-bias, such as a tendency not to repeat recently unrewarded decisions (Abrahamyan et al. 2016), are likely to contribute substantially in a decision-making system, but are unrelated to the statistics of the stimulus and as such can not be captured by this model. To summarize, we uncovered history effects in the model that are due to statistical changes in the observed samples. The overall statistical structure adapts on a long timescale, while small biases arise on a short time scale.

**Fig. 5.**
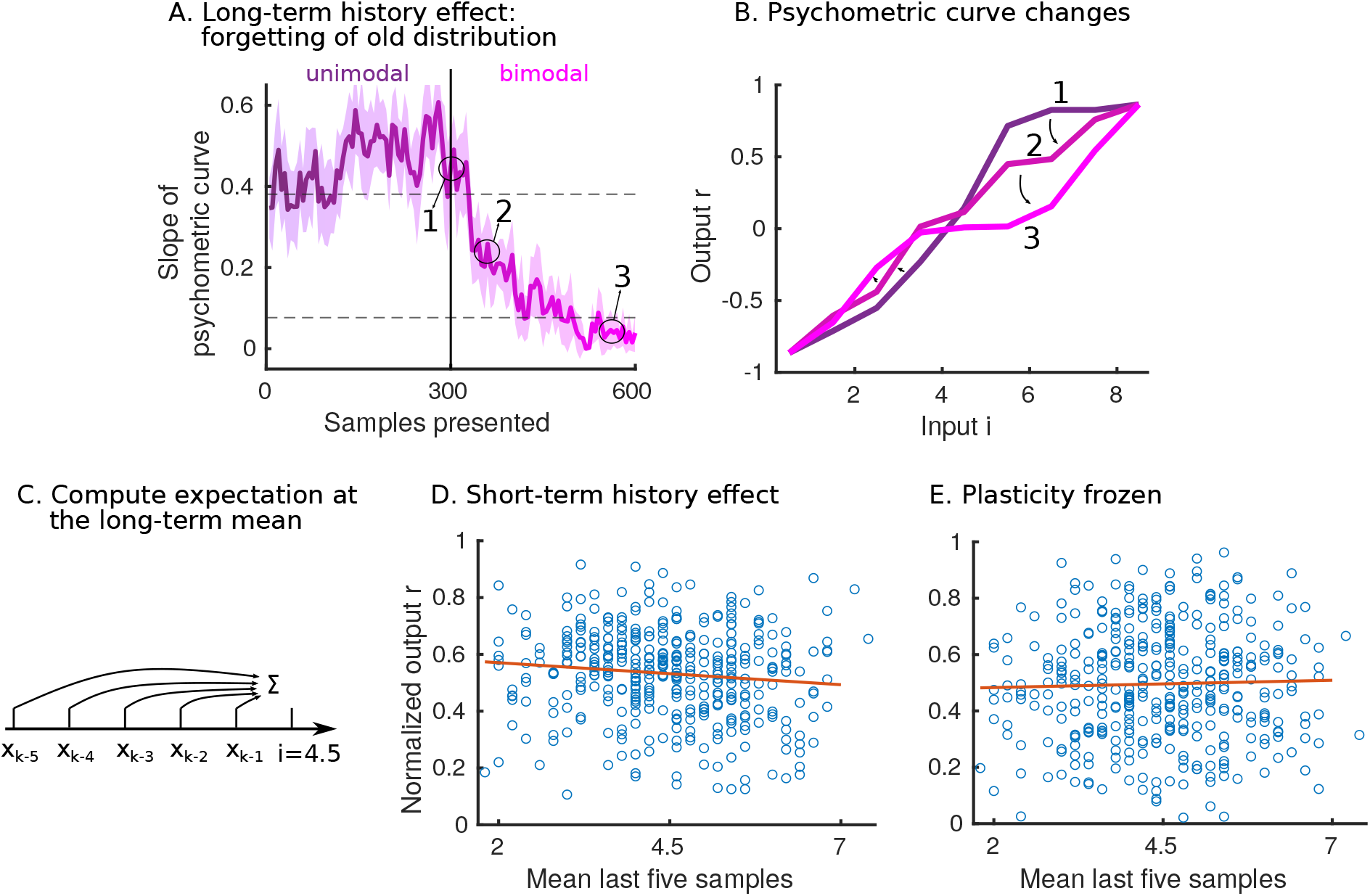
The model exhibits long- and short-term history effects. (A) The target distribution presented for the first 300 samples is unimodal (as in Fig 4C). The target distribution presented for the last 300 samples is bimodal (as in Fig 4C). The slope of the psychometric curve is measured every fifth sample. The shaded area indicates the standard deviation computed from 10 simulations. The horizontal dashed lines indicate the slope of the psychometric curves of the unimodal (top) and bimodal (bottom) target distributions. (B) The psychometric curves at three time points, indicated by arrows in panel (A), are shown (averaged over 20 simulations). (C) Cartoon of simulation. After every fifth sample, we simulate a decision using the input *i* = 4.5, which is the long-term mean of the target distribution. There is no switch between target distributions, we are at steady-state and providing samples from the bimodal distribution only (as in Fig 4C). We look at the effect of the last five samples on the output *r*. (D) The blue circles indicate simulation results. The output *r* is normalized to the interval [0,1] and plotted as a function of the mean of the last five samples. The red line is a result of linear regression, the slope is significantly non-zero (p-value 0.0257). (E) The same plot as in (D), but plasticity is frozen. The slope is not significantly non-zero (p-value 0.56).

### The model can recall the probabilities instantaneously

The model produces samples, according to the inverse transform stored into the weights from the uniform sampling network to the sensory network. The estimation of the probabilities is therefore only accurate after waiting for a few seconds (Fig 3). This representation can, however, be transformed into a different, more instantaneous representation. We provide an example of how to encode the probability of a stimulus directly in the activity of a read-out network. First, a read-out network is connected to the sensory network (Fig 6A). This read-out network is balanced, consisting of one pool of excitatory neurons and one pool of inhibitory neurons. All excitatory neurons from the sensory network connect to all excitatory neurons of the read-out network. These weights follow a short-term plasticity (STP) rule. Specifically, the weights depress when the presynaptic neuron is active (see Methods). This means that read-out weights depress more when a cluster of excitatory neurons in the sensory network is more active, leading to lower activity in the read-out network. This directly corresponds to the probability of the stimulus for which the cluster codes, i.e. there is an inverse relationship between the network activity of the read-out and the probability of the stimulus. This can be interpreted as a novelty signal, where low-probability stimuli lead to high activity and vice versa. This relationship between read-out network activity and input probability need not be linear for the entire range of probabilities, as varying STP parameters give different activity profiles (Fig S4A). When synapses facilitate instead of depress, we see the opposite behaviour: the network activity of the read-out monotonously increases with the probability of the stimulus (Fig S4B). We conclude that the sampling representation can be accessed in different ways. Not only can we compute expectations over functions, but we can also transform the representation and use it for instantaneous coding.

**Fig. 6.**
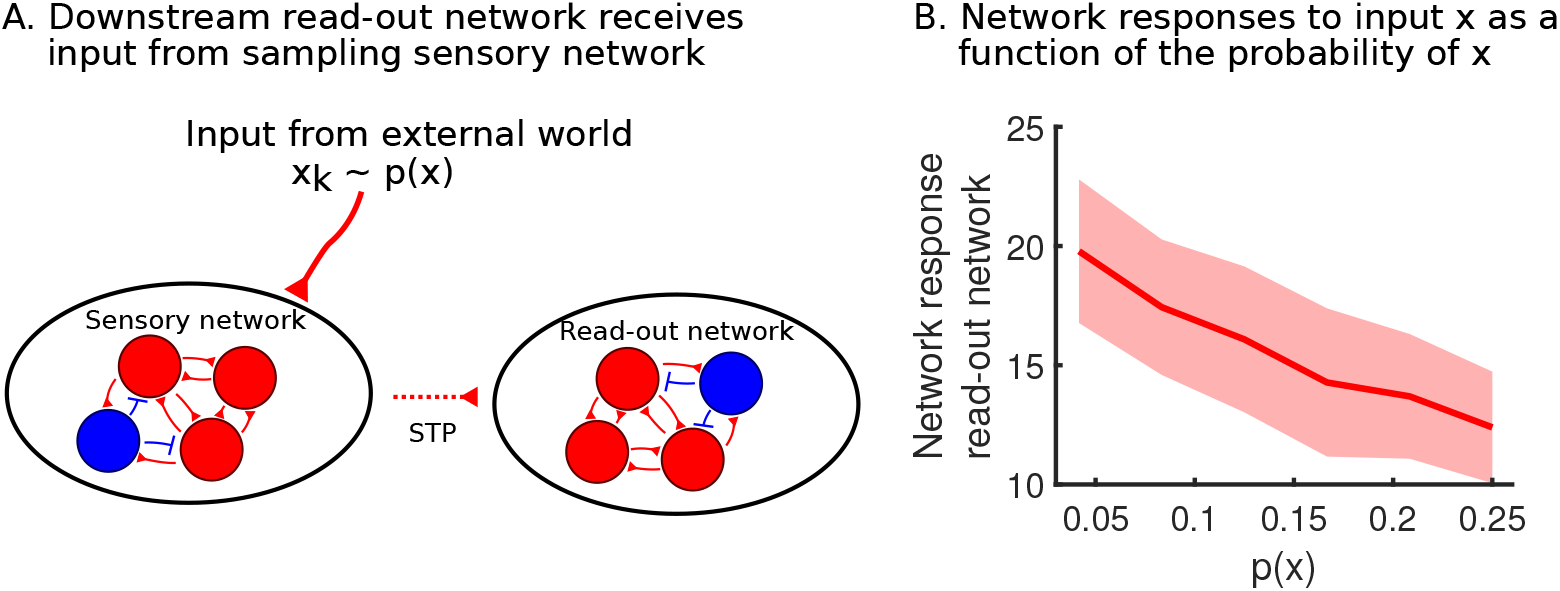
The model can recall the probabilities instantaneously. (A) A read-out network with STP in the read-out weights can encode probabilities instantaneously. (B) There is an inverse relationship between probability and average read-out activity when using short-term depression in the read-out weights. Shaded area indicates one standard deviation from the mean on each side (25 simulations).

## DISCUSSION

### Summary

We presented a model which learns the probability distribution of an external variable. The model takes samples from the distribution during spontaneous dynamics, using inverse transform sampling. This representation can compute expectations over functions in downstream networks. Specifically, we studied a possible relationship with perceptual decision-making. The model predicts that the shape of the psychometric curves depends on the stimulus distribution. Additionally, sensory history effects emerge due to the ongoing plastic changes. Finally, we explored a way in which the representation can be transformed: it is possible to transform the samples into an instantaneous coding of the probability of the external variable.

### Learning with clustered networks

The substrate we used for learning statistical structure is the clustered network. One clustered network serves as a backbone and projects to a second network which encodes the variable. The plastic weights learn the correct transformation, in this case, the inverse of the cumulative distribution function *F*^−1^ (*u*). This is related to previous work on learning and generating sequences (Nicola and Clopath 2017; Maes et al. 2020; Maes et al. 2021). In this previous work, the backbone consists not of randomly switching clusters but clusters active sequentially in a chain. Instead of encoding the uniform distribution, it encodes time. The plastic weights to the network encoding the variable learn a function of time *f*(*t*), using similar plasticity rules. In both models, the architecture is identical, leading to comparable design features: for example, the accuracy and speed of learning depend to a large extent on the number of clusters in the backbone. A clustered connectivity has also been shown to enhance reinforcement learning in a recent study (Weidel et al. 2021). More generally, a clustered code has interesting properties, such as robust error correction, that make it a candidate to underlie computations in the brain (Berry and Tkačik 2020).

### Modularity leads to flexibility

A strength of the model is its large flexibility, stemming from its modularity (Koblinger et al. 2021). Once the statistical structure is stored, it can be accessed in several ways. Separating the storage of the statistical structure from performing downstream computations is also observed in the experimental literature in the context of working-memory tasks (Akrami et al. 2018; Loewenstein et al. 2021). In particular, when the downstream computation does not change, but the statistical structure does change, it may be sufficient to have unsupervised learning to update the stored distribution. When the computation itself changes, a form of reward-based or supervised learning could act in the downstream networks leaving the statistical structure unchanged.

### Computing expectations and transforming the representation

We focused on a model of sampling in spiking neural networks. We illustrated the use of the representation in downstream networks in two examples. Many different mechanisms could work complementary to our model, both when performing decision-making and when encoding surprise or novelty in a network. The generated samples might be one of many inputs to a hierarchical decision-making system, relying on more than temporal integration (Sarafyazd and Jazayeri 2019; Cowley et al. 2020; Tervo et al. 2021). Furthermore, novelty signals have been theorized to emerge from various sources. Previous studies have proposed, similarly, plasticity mechanisms in feedforward excitatory synapses (Park and Geffen 2020; Maoz et al. 2020). Other mechanisms however are also likely to be involved, for example, inhibitory plasticity onto excitatory neurons could suppress non-novel stimuli (Ramaswami 2014; Schulz et al. 2020). More work has to be done to reveal all the mechanisms underlying this phenomenon and how they interact together.

### Other types of statistical structure

Our work focuses on one type of statistical structure: a prior probability distribution. The brain may extract other types of structure to influence behaviour. One other such type is Markov statistics. Certain events may precede other events with a high or low probability, potentially informing our predictions and decisions. A conceptual model was proposed before, but its implementation in a neural system remains non-trivial (Bernstein et al. 2017). Finally, an open question is how the stored statistical structure can be integrated with working memory for more complex decision-making. This remains to be explored in future work.

### Other types of sampling

Interpreting neural activity as samples from a distribution is not new in itself. Many different studies have investigated this idea (Hoyer and Hyvärinen 2003; Fiser et al. 2010; Berkes et al. 2011; Orbán et al. 2016; Haefner et al. 2016) and have made progress towards implementing sampling into spiking networks (Buesing et al. 2011; Moreno-Bote et al. 2011; Savin and Deneve 2014). Our work shows how to implement a well-known mechanism, inverse transform sampling, in a biophysically realistic network. Additionally, we show how it is possible to learn from observations, as opposed to the previously hard-coded implementations. Inverse transform sampling is a form of direct sampling, where there are no autocorrelations between the samples since the uniform samples are independently drawn. This is in contrast to sampling techniques such as Markov Chain Monte Carlo (MCMC), where correlations between subsequent samples are unavoidable during the stochastic walk in the probability landscape. Our proposed way of sampling is particularly useful when the distributions have a low dimensionality. But, when the dimensionality increases and the curse strikes, MCMC algorithms become beneficial. Previous work has focused on versions of MCMC, in the context of sampling from high-dimensional posterior distributions (Savin and Deneve 2014; Zhang et al. 2020b; Zhang et al. 2020a). This makes sense especially when investigating sensory processing, where a high-dimensional input, such as an image, has to be processed in noisy circumstances. Another recent study in this context implemented a distinct type of sampling by optimizing the recurrent connectivity of a network, minimizing a cost function (Echeveste et al. 2020). We focus here, however, on external variables that are more salient and cognitively relevant, and, as such, are of much lower dimensionality. Yet other work has explored mental sampling in the context of foraging and free-recall experiments (Zhu et al. 2018), proposing more complex hierarchical sampling than is done in either our model or standard MCMC.

### Conclusion

We studied a model capable of learning to sample from a target prior distribution by mapping inverse transform sampling into a spiking network architecture. We propose that the sampling representation can serve as a basis for downstream computations, and provide testable predictions in the case of perceptual decision-making.

## METHODS AND MATERIALS

Excitatory neurons (*E*) are modelled with the adaptive exponential integrate-and-fire model (Brette and Gerstner 2005). A classical integrate-and-fire model is used for the inhibitory neurons (*I*).

### Model architecture

#### Sensory network

The sensory network encodes the external variable, and consists of 8 clusters of 100 excitatory neurons and a pool of 200 inhibitory neurons. The connection strengths in the sensory network are found in Table 1, with a scaling factor *f* = 1. These connection strengths roughly correspond to values found in Litwin-Kumar and Doiron 2014; Maes et al. 2020, where the *E* to *E* and *I* to *E* synapses are plastic. There is no all-to-all connectivity; rather two neurons are connected with a probability of *p*. The synaptic connections within the same cluster are multiplied by a factor of 10.

**TABLE 1.**
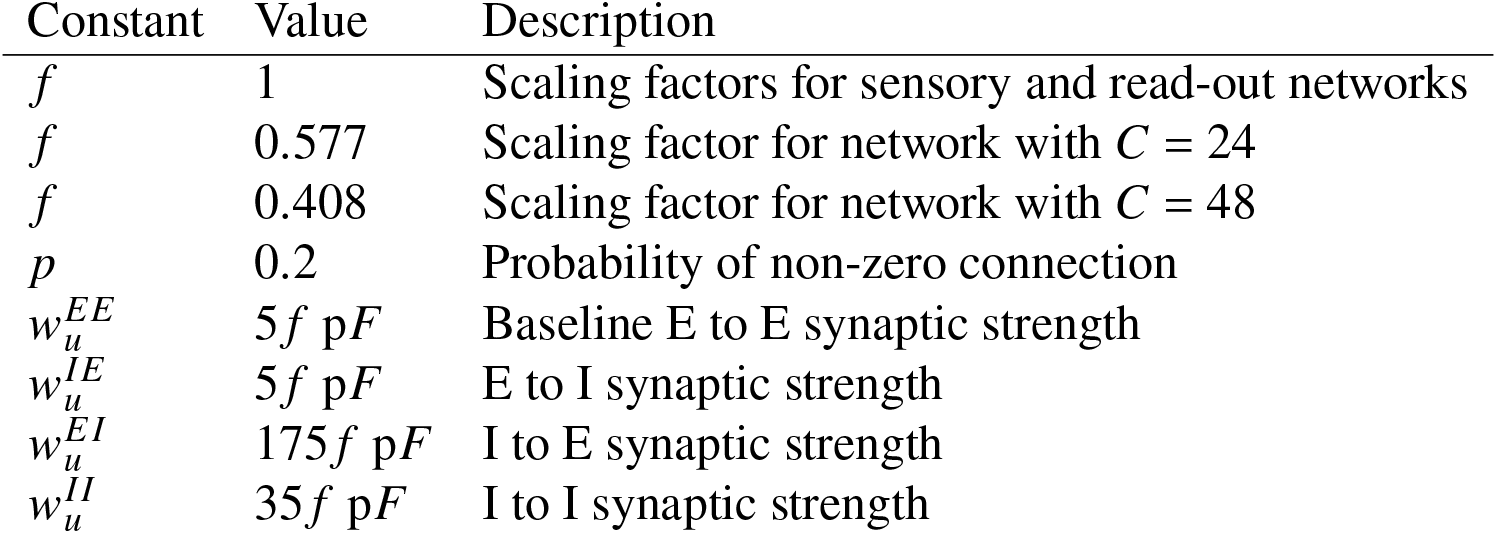
Network parameters.

#### Uniform sampler network

The uniform sampler network is a network which spontaneously switches activity between *C* clusters of excitatory neurons. Each cluster consists of 100 excitatory neurons and there is a pool of 25C inhibitory neurons. In our study, we simulate two uniform sampler networks of a different size. We use the network parameters of the smaller sensory network and scale those parameters by a scaling factor *f*, proportional to the square root of the relative network sizes. The synaptic connections within the same cluster are multiplied by a factor of 20, for the network with *C* = 24, and a factor of 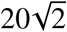, for the network with *C* = 48. Network connectivities are found in Table 1.

#### Read-out network

The read-out network receives input from the sensory network. It consists of 800 excitatory neurons and 200 inhibitory neurons, all the excitatory neurons receive input from all the excitatory neurons in the sensory network. The connection strengths in the network are the same as the connections strengths in Table 1, with *f* = 1.

### Neural and synaptic dynamics

All neurons in the model are either excitatory (*E*) or inhibitory (*I*). The parameters of the neurons do not change depending on which network they belong to. Parameters are taken from Brette and Gerstner 2005; Clopath et al. 2010; Litwin-Kumar and Doiron 2014; Maes et al. 2020; Maes et al. 2021.

#### Membrane potential dynamics

The membrane potential of the excitatory neurons (*V^E^*) has the following dynamics:

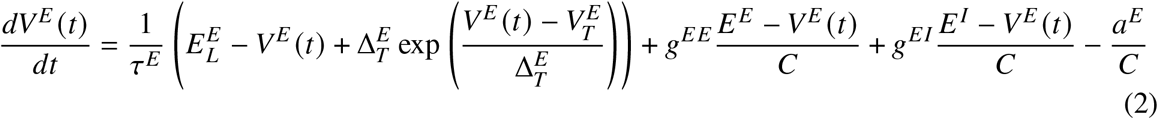

where *τ^E^* is the membrane time constant, 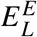 is the reversal potential, 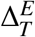 is the slope of the exponential, *C* is the capacitance, *g^EE^, g^EI^* are synaptic inputs from excitatory and inhibitory neurons respectively and *E^E^, E^I^* are the excitatory and inhibitory reversal potentials respectively. When the membrane potential exceeds 20 mV, the neuron fires a spike and the membrane potential is reset to *V_r_*. This reset potential is the same for all neurons in the model. There is an absolute refractory period of *τ_abs_*. The parameter 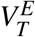 is adaptive for excitatory neurons and set to *V_T_* + *A_T_* after a spike, relaxing back to *V_T_* with time constant *τ_T_*:

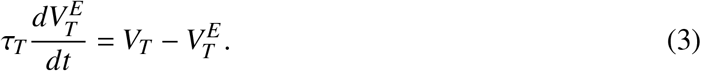

The adaptation current *a^E^* for excitatory neurons follows:

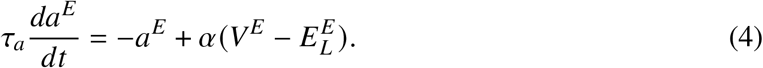

where *τ_a_* is the time constant for the adaptation current. The adaptation current is increased with a constant *β* when the neuron spikes. The constant *β* is larger in the uniform sampler network. This makes the switching dynamics less random in time, i.e. the switching happens reliably at about 8 Hz.

The membrane potential of the inhibitory neurons (*V^I^*) has the following dynamics:

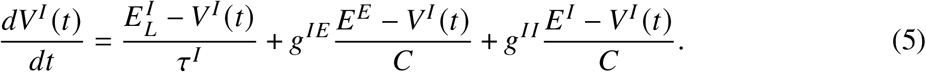

where *τ^I^* is the inhibitory membrane time constant, 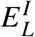 is the inhibitory reversal potential and *E^E^, E^I^* are the excitatory and inhibitory resting potentials respectively. *g^EE^* and *g^EI^* are synaptic input from excitatory and inhibitory neurons respectively. Inhibitory neurons spike when the membrane potential crosses the threshold *V_T_*, which is non-adaptive. After this, there is an absolute refractory period of *τ_abs_*. There is no adaptation current (see Table 2 for the parameters of the membrane dynamics).

**TABLE 2.**
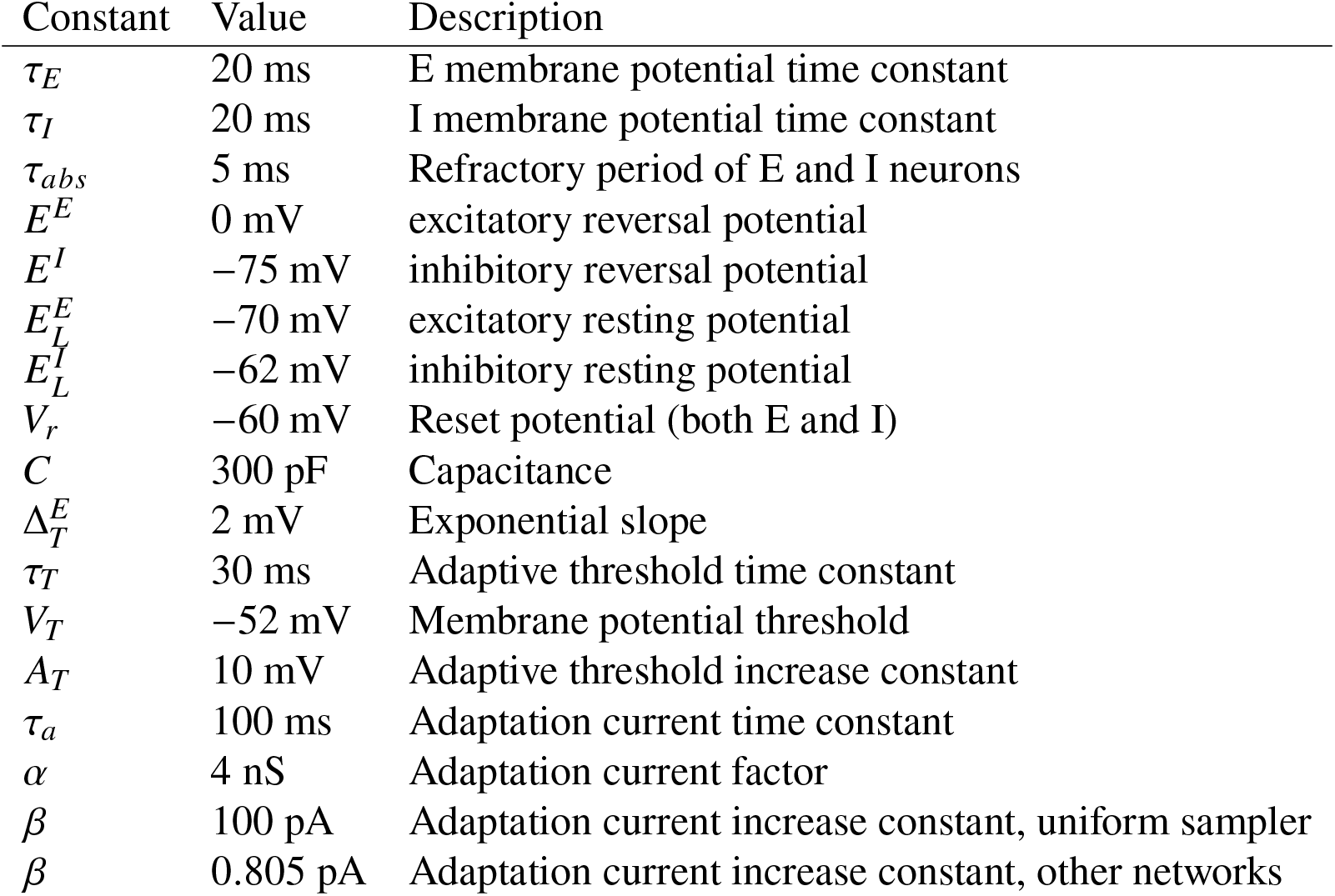
Neuronal membrane dynamics parameters. Parameters taken from Brette and Gerstner 2005; Clopath et al. 2010; Litwin-Kumar and Doiron 2014; Maes et al. 2020; Maes et al. 2021.

| Constant | Value | Description |
| --- | --- | --- |
| $\tau_E$ | 20 ms | E membrane potential time constant |
| $\tau_I$ | 20 ms | I membrane potential time constant |
| $\tau_{abs}$ | 5 ms | Refractory period of E and I neurons |
| $E^E$ | 0 mV | excitatory reversal potential |
| $E^I$ | -75 mV | inhibitory reversal potential |
| $E_L^E$ | -70 mV | excitatory resting potential |
| $E_L^I$ | -62 mV | inhibitory resting potential |
| $V_r$ | -60 mV | Reset potential (both E and I) |
| $C$ | 300 pF | Capacitance |
| $\Delta_T^E$ | 2 mV | Exponential slope |
| $\tau_T$ | 30 ms | Adaptive threshold time constant |
| $V_T$ | -52 mV | Membrane potential threshold |
| $A_T$ | 10 mV | Adaptive threshold increase constant |
| $\tau_a$ | 100 ms | Adaptation current time constant |
| $\alpha$ | 4 nS | Adaptation current factor |
| $\beta$ | 100 pA | Adaptation current increase constant, uniform sampler |
| $\beta$ | 0.805 pA | Adaptation current increase constant, other networks |

#### Synaptic dynamics

The synaptic conductance, *g*, of a neuron *i* is time dependent, it is a convolution of a kernel with the total input to the neuron *i*:

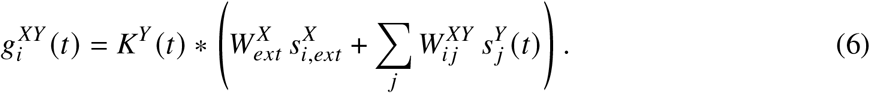

where *X* and *Y* can be either *E* or *I*. 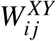 is the synaptic strength from presynaptic neuron *j* to postsynaptic neuron *i*, and 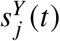 is one when the presynaptic neuron *j* spikes and zero otherwise.

*K* is the difference of exponentials kernel:

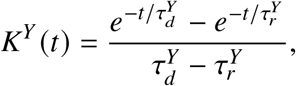

with a decay time *τ_d_* and a rise time *τ_r_* dependent only on whether the neuron is excitatory or inhibitory. The conductance is a sum of recurrent input and external input. The externally incoming spike trains 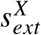 are generated from a Poisson process with rates 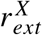. The excitatory external input to the uniform sampler network depends on the number of clusters, tuned to give a similar rate of switching between the clusters. The excitatory external input to the sensory network is slightly lower because it also receives excitatory input from the uniform sampler. The externally generated spike trains enter the network through synapses 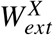. Parameters for the synaptic dynamics are found in Table 3. Parameters were not fine tuned. They are set to match similar activities across the different networks, and are taken from Litwin-Kumar and Doiron 2014; Maes et al. 2020; Maes et al. 2021.

**TABLE 3.** Synaptic dynamics parameters. Parameters taken from Litwin-Kumar and Doiron 2014; Maes et al. 2020; Maes et al. 2021.

| Constant | Value | Description |
| --- | --- | --- |
| $\tau_d^E$ | 6 ms | E decay time constant |
| $\tau_r^E$ | 1 ms | E rise time constant |
| $\tau_d^I$ | 2 ms | I rise time constant |
| $\tau_r^I$ | 0.5 ms | I rise time constant |
| $W_{ext}^E$ | 1.6 pF | External input synaptic strength to E neurons |
| $r_{ext}^E$ | 5.5 kHz | Rate of external input to E neurons of uniform sampler ($C = 48$) |
| $r_{ext}^E$ | 5 kHz | Rate of external input to E neurons of uniform sampler ($C = 24$) |
| $r_{ext}^E$ | 4 kHz | Rate of external input to E neurons of sensory network |
| $W_{ext}^I$ | 1.52 pF | External input synaptic strength to I neurons |
| $r_{ext}^I$ | 2.25 kHz | Rate of external input to I neurons |

### Plasticity

The synaptic weight from excitatory neuron *j* in the uniform sampler network to excitatory neuron *i* in the sensory network is changed according to the following differential equation:

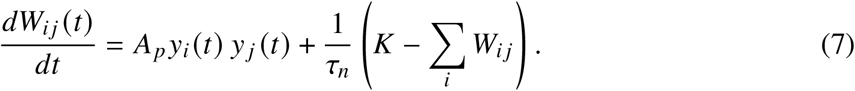

where *y_i_*(*t*) = 1 if the postsynaptic neuron *i* fired in the last 15 ms and zero else. Similarly for the presynaptic neuron *y_j_*(*t*) = 1 if the presynaptic neuron *j* fired in the last 15 ms and zero else. *A_p_* is the amplitude of synaptic potentiation. The second term is a “soft” normalization. The normalization ensures that probability mass can be smoothly re-attributed, specifically it ensures that each cluster of neurons in the uniform sampler network connects to a single cluster in the sensory network. *τ_n_* is the time constant of the normalization and *K* the normalization constant. Weights vary between [*W_min_, W_max_*]. Parameters are found in Table 4.

**TABLE 4.**
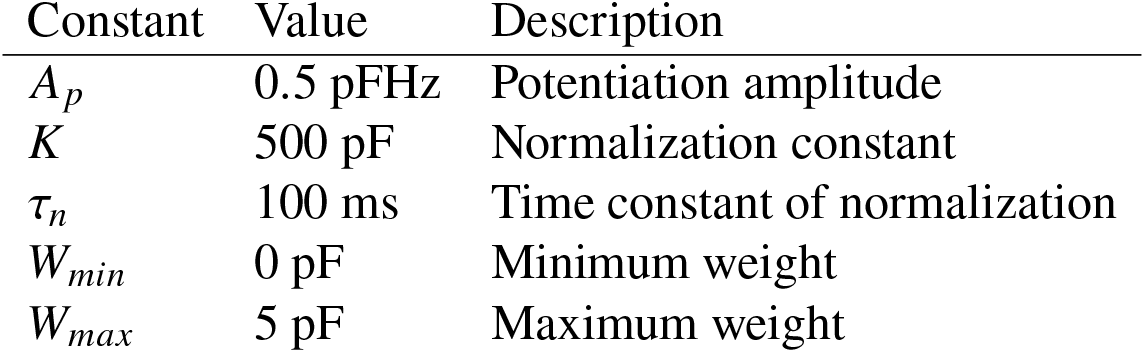
Plasticity parameters.

### Numerical simulations

#### Protocol

During learning, samples from the target distribution are drawn *x_k_* ~ *p*(*x*) every 200 ms. A high external input (30 kHz) is given to the cluster *k* of excitatory neurons corresponding to sample *x_k_*, for 50 ms. During spontaneous activity, the baseline external input is given (Table 3).

#### Learning curve

Learning curves can be obtained using the weights from the uniform sampler network to the sensory network. All the weights to cluster *k* of the sensory network are summed and divided by the total sum of all the plastic weights. This gives an empirical distribution, which can directly be compared with the target distribution. The weights were saved every fifth sample presentation. The MATLAB function *ranksum* is used to perform the Mann-Whitney U-test in Fig 2D.

#### KL-divergence

The KL-divergence is a measure of distance between two probability distributions. Consider the spike trains in the sensory network until time *t*. At each moment in time, only one of the clusters is active. The active cluster is determined by convolving the spike trains with a Gaussian of width 20 ms, averaging over the clusters, and taking the maximum. The amount of time that each cluster is active, divided by the total time *t* is the empirical probability that a cluster is active. Denote 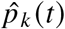 as the empirical probability of cluster *k* at time *t*. The KL-divergence is then:

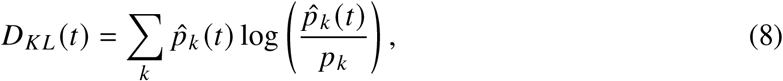

where *p_k_* is the target probability for cluster *k*. The KL-divergence decreases with time, indicating a better match between empirical and target distributions as more samples are accumulated. The dashed lines in Fig 3B are computed in the exact same way. The difference is that the samples are not obtained from the neural activity in the sensory network, but by using the random number generator of MATLAB (built-in function *rand*).

#### Computing expectations

Nine inputs *i* are given, *i* = 0.5,…, 8.5. To compute *f*(*i, x_t_*), samples *x_t_* are obtained in continuous time by running the spontaneous dynamics of the model. At each time *t*, the neural activity in the sensory network is averaged over clusters. The index of the cluster which is the most active at time *t* gives *x_t_*. For example, if cluster 3 is the most active at time *t* we have *x_t_* = 3. The output *r* integrates the function with a time constant *τ_r_* = 1000 ms. We assume the eventual decision to be a function of the output *r*. However, we do not implement the actual decision in a neural circuit as we are agnostic about the precise way the evaluation and integration of *f* (*i, x_t_*) happens. Mechanistic implementations have been proposed before, for example using attractor network models (Wong and Wang 2006; Esnaola-Acebes et al. 2021).

Slopes of psychometric curves are computed for figures 4E and S3A. The slopes are computed by saving the outputs *r* when given input *i* = 5.5 and input *i* = 3.5. The resulting outputs *r* are subtracted and divided by two. In figure 4E, the slope is also normalized by the output *r* when given an input *i* = 8.5; this normalization is important to be able to compare the slopes for varying simulation time lengths. The MATLAB function *fitlm* is used to fit linear regression and compute the p-value for the short-term history effect in Figs 5D-E.

#### Short-term plasticity

Short-term plasticity is implemented for the instantaneous decoding of probability (see Fig 6). All excitatory neurons in the sensory network are connected to all excitatory neurons in the read-out network. For all these read-out weights, we have a baseline connectivity strength of *w* = 4 pF. Weights *w_j_* from neuron *j* in the sensory network to all neurons in the read-out network are depressed when neuron *j* fires, by an amount of 0.05*w_j_*, bounded at zero. Depressed weights return exponentially back to baseline strength with a time constant of 2 s. The same constants are used for the simulation using facilitation (see Fig S4B). The strength of the read-out weight increases by an amount of 0.05*w_j_* at every presynaptic spike (maximum is *w* = 6 pF) and decays to zero with a time constant of 2 s.

#### Simulations

The code used for the training and testing of the spiking network model is built in Matlab. Forward Euler discretisation with a time step of Δ*t* = 0.1 ms is used. The code will be made available after publication.

## ACKNOWLEDGMENTS

AM acknowledges funding through the EPSRC Centre for Neurotechnology. MB acknowledges funding through EPSRC award EP/N014529/1 supporting the EPSRC Centre for Mathematics of Precision Healthcare at Imperial. CC acknowledges support by BBSRC BB/N013956/1, BB/N019008/1, Wellcome Trust 200790/Z/16/Z, Simons Foundation 564408 and EPSRC EP/R035806/1.

## SUPPLEMENTARY FIGURES

**Fig. S1.**
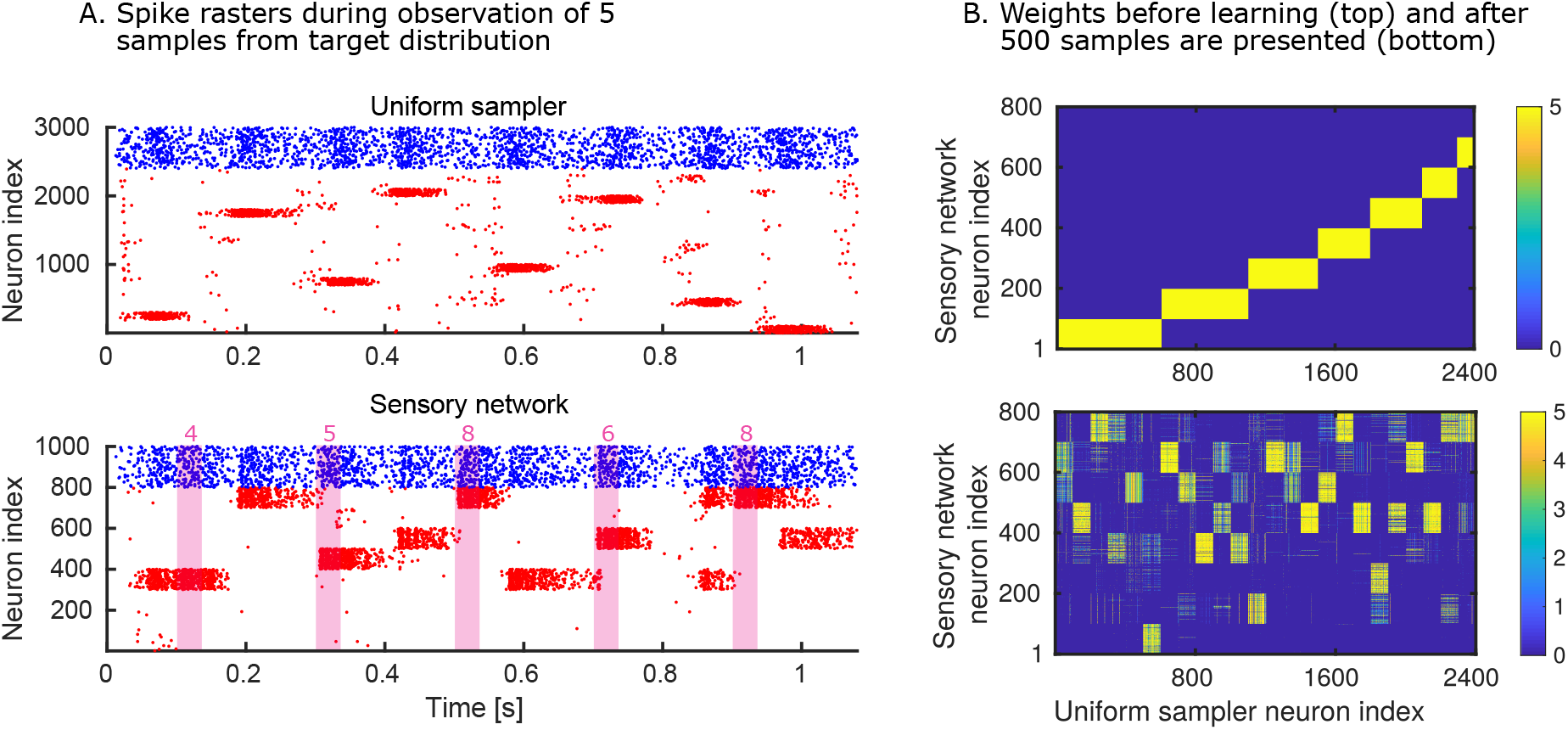
Activity during sequential observations and weight changes. (A) Spike rasters of the uniform sampler network (top) and the sensory network (bottom) during the sequential observation of five samples from the target distribution. External input is given to clusters in the sensory network during the observation of a sample (for 50 ms, indicated in pink shading). The numbers in pink are the samples and indicate which cluster is stimulated. The activity in the uniform sampler does not need to be synchronized with the observation of stimuli. (B) Weights from the uniform sampler network to the sensory network. The weights are initialized to encode the initial distribution in Fig 2B (top). The weights change by observing samples (bottom).

**Fig. S2.**
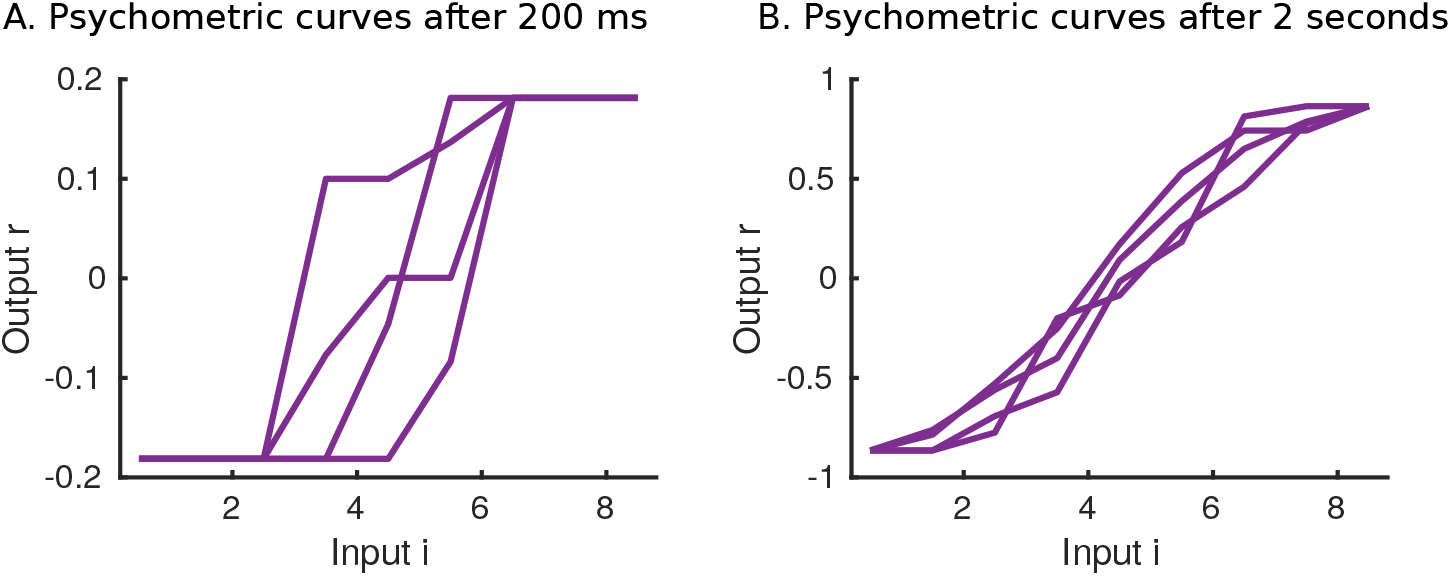
Individual psychometric curves. (A) We plot 4 psychometric curves, resulting from 4 repetitions of a 200 ms decision. (B) We plot 4 psychometric curves, resulting from 4 repetitions of a 2 seconds decision. Longer decision times lead to less variability between individual curves. Psychometric curves shown in the main text are averaged over many such individual curves.

**Fig. S3.**
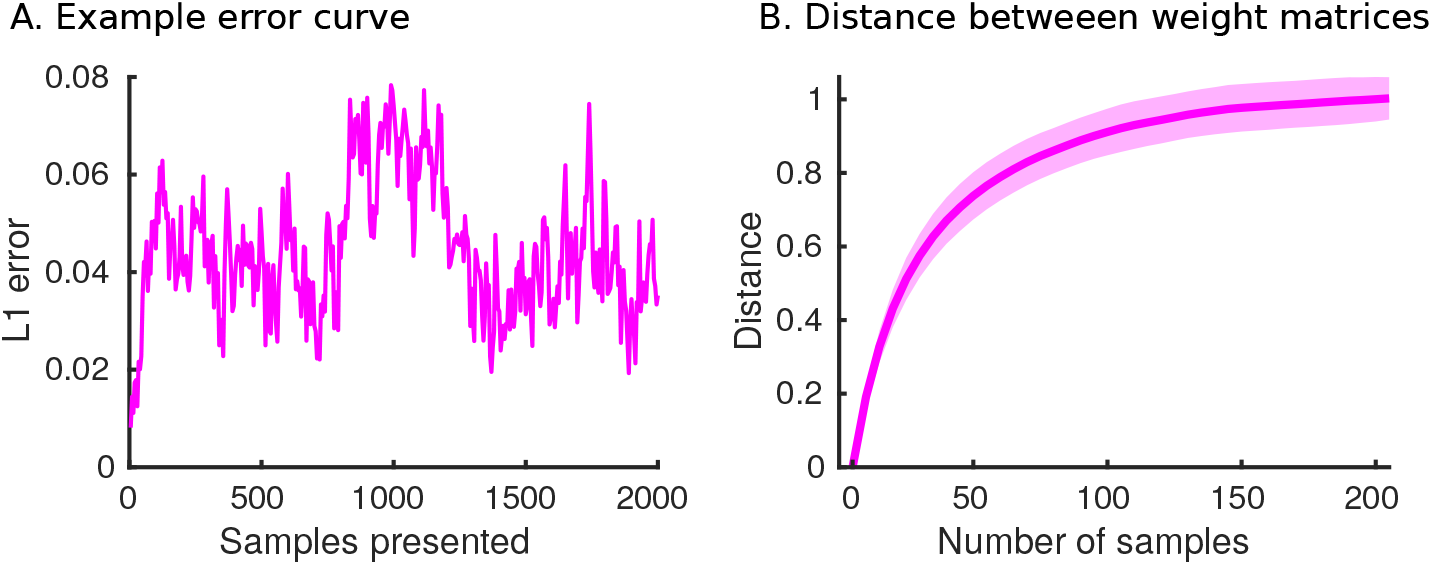
History effects due to ongoing rewiring. (A) The error between the distribution stored by the model and the target distribution (here: bimodal distribution) fluctuates even when the target distribution is kept constant. This is due to constant rewiring of the plastic weights. (B) We record the plastic weight matrices at different time points. The distance between two plastic weight matrices increases with the amount of samples presented between their recording times. The distance is measured by treating the matrices as vectors, and taking the L1 norm of their difference. The weights from the simulation in panel (A) are used here. The shaded area indicates the standard deviation computed from 360 simulations.

**Fig. S4.**
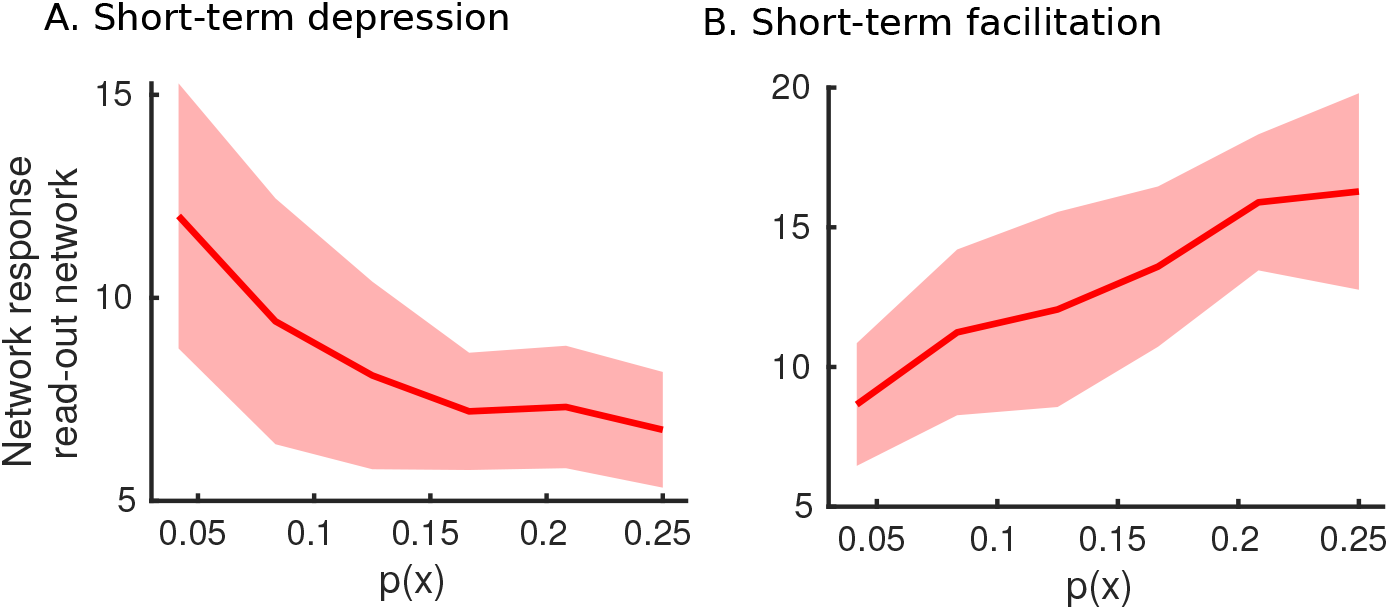
The model can instantaneously encode probabilities with both short-term depression and facilitation. (A) Linear relationship at low probabilities (*p*(*x*) < 0.15) and near constant at higher probabilities (*p*(*x*) > 0.15). The depression parameter was increased four-fold (compared to Fig 6B), the time constant was kept the same. (B) Using facilitation will lead to the amplification of high-probability inputs (to be compared with Fig 6B). The shaded area indicates one standard deviation from the mean in both panels (A) and (B), measured over 25 simulations.

